# Sex differences in the ability of corticostriatal oscillations to predict rodent alcohol consumption

**DOI:** 10.1101/750711

**Authors:** Angela M. Henricks, Emily D.K. Sullivan, Lucas L. Dwiel, Karina M. Keus, Ethan D. Adner, Alan I. Green, Wilder T. Doucette

## Abstract

**Background:** Although male and female rats differ in their patterns of alcohol use, little is known regarding the neural circuit activity that underlies these differences in behavior. The current study used a machine learning approach to characterize sex differences in local field potential (LFP) oscillations that may relate to sex differences in alcohol drinking behavior.

**Methods:** LFP oscillations were recorded from the nucleus accumbens shell and the rodent medial prefrontal cortex of adult male and female Sprague-Dawley rats. Recordings occurred before rats were exposed to alcohol (n=10/sex X 2 recordings/rat) and during sessions of limited access to alcohol (n=5/sex X 5 recordings/rat). Oscillations were also recorded from each female rat in each phase of estrous prior to alcohol exposure. Using machine-learning, we built predictive models to classify rats based on: 1) biological sex; 2) phase of estrous; and 3) alcohol intake levels. We evaluated model performance from real data by comparing it to the performance of models built and tested on permutations of the data.

**Results:** Our data demonstrate that corticostriatal oscillations were able to predict alcohol intake levels in males (p<0.01), but not in females (p=0.45). The accuracies of models predicting biological sex and phase of estrous were related to fluctuations observed in alcohol drinking levels; females in diestrus drank more alcohol than males (p=0.052), and the male vs. diestrus female model had the highest accuracy (71.01%) compared to chance estimates.

Conversely, females in estrus drank similar amounts of alcohol to males (p=0.702), and the male vs. estrus female model had the lowest accuracy (56.14%) compared to chance estimates.

**Conclusions:** The current data demonstrate that oscillations recorded from corticostriatal circuits contain significant information regarding alcohol drinking in males, but not alcohol drinking in females. Future work will focus on identifying where to record LFP oscillations in order to predict alcohol drinking in females, which may help elucidate sex-specific neural targets for future therapeutic development.

## Background

Alcohol use contributes to 5.1% of the global disease burden, accounting for 5% of all deaths in men and 1% of all deaths in women in the United States alone (1–3). While historically men drink more alcohol than women, this gender gap is closing (4), and women tend to escalate to alcohol dependence more rapidly than men (2,5). Though these sex differences partly arise from sociocultural factors, there are known sex differences in the activity of brain regions that underlie substance use behavior (5,6). However, the specific neurobiological underpinnings contributing to sex differences in alcohol drinking are poorly understood, limiting the development of more efficacious, targeted therapies for problematic alcohol use.

One barrier to the development of better therapies for excessive alcohol use is the fact that the majority of preclinical neuroscience studies have used only male animals (7,8). However, the available behavioral data in rodent models of alcohol drinking demonstrate that female rats, in a non-dependent state, drink more alcohol and show greater alcohol preference than male rats (9), as well as display heightened sensitivity to the rewarding effects of alcohol compared to males (10). The behavioral differences between females and males are biological in nature as neonatal masculinization of females reduces alcohol intake compared with intact female rats, resulting in patterns of drinking similar to those displayed by males (11). In a similar study, intact female rats showed a heightened reward response to alcohol than either males or ovariectomized females, suggesting that ovarian hormones help facilitate the reinforcing properties of alcohol (10). Ovarian hormone status has also been associated with small fluctuations in alcohol consumption in intact females (12,13). However, it is currently unknown whether the neural circuits that regulate alcohol consumption show sexually dimorphic activity patterns (and whether these patterns are influenced by ovarian hormone status) that may explain the sex differences in alcohol drinking behavior.

The mechanistic role of corticostriatal circuits in regulating the rewarding properties of alcohol is well characterized in male rodents (14). In rats (and humans), the nucleus accumbens (NAc) integrates cortical inputs and indirectly sends feedback to frontal brain regions (medial prefrontal cortex in humans [mPFC]; prelimbic [PL] and infralimbic [IL] cortices in rats) (15), and is particularly important in the motivating properties of abused drugs (16). The mPFC is also activated in response to reward-related cues, and it has been suggested that deficits in the ability to inhibit responses to drugs arises from dysregulated communication between the mPFC and striatal regions (17). Thus, we hypothesize that male and female rats might display inherent (i.e., trait-level) differences in corticostriatal circuit activity, which may be associated with sex differences in alcohol drinking behaviors.

Activity in the corticostriatal circuit can be examined longitudinally by measuring local field potential (LFP) oscillations in awake, freely behaving rats. LFP oscillations provide a readout of electrical potential from a group of neurons that relates to individual neuronal activity, as demonstrated by neuronal phase locking and ensemble classification (18–20). LFP oscillations recorded from reward-related regions have been shown to change during behavior (21) and reflect pharmacologic manipulation (22–24). For instance, in male rats, low frequency oscillations decrease while high frequency oscillations increase following an injection of alcohol (25). Furthermore, low frequency oscillations in the cortex and NAc appear to be hypoconnected in alcohol preferring rats (sex not reported) compared to outbred rats, which was reversed by alcohol exposure (26). LFP oscillations can therefore be a valuable readout of circuit dynamics related to alcohol drinking behaviors (i.e., amount of alcohol consumed) in rodents.

In the current experiment, we measured corticostriatal LFP oscillations in adult male and female rats prior to and during alcohol drinking behavior. Using an unbiased machine learning approach, we aimed to determine whether LFPs recorded from corticostriatal circuits contained information regarding: 1) biological sex; 2) ovarian hormone status; and 3) the amount of alcohol consumed during an alcohol drinking session. We hypothesized that sex differences in inherent corticostriatal circuit activity might be related to sex differences in alcohol drinking behavior.

## Methods

### Subjects and Housing

Male and female Sprague-Dawley rats (n = 10/sex) were purchased from Charles River (Wilmington, MA, USA) and arrived on postnatal day 60. All animals were housed individually on a reverse 12-hour light cycle with *ad libitum* access to food and water. All experiments were carried out in accordance with the National Institute of Health Guide for the Care and Use of Laboratory Animals (NIH Publications No. 80-23) and were approved by the Institutional Animal Care and Use Committee of Dartmouth College.

### Electrode Construction and Implantation

Electrodes were designed and constructed in-house and were similar to those used in our previous publication (27). Animals were anesthetized with isoflurane gas (4% induction, 2% maintenance) and secured into a stereotaxic frame. Custom electrodes were implanted bilaterally targeting the NAc shell (NAcSh; from bregma: DV −8mm; AP +1.2mm; ML +/−1.0mm) and PL/IL junction of the mPFC (from bregma: DV −5mm; AP +3.7mm; ML +/−0.75mm). Four stainless steel skull screws were placed around the electrode site and dental cement (Dentsply, York, PA, USA) was applied to secure the electrodes in place.

### Recording and Processing Local Field Potential Oscillations

LFP oscillations were recorded in sound-attenuated chambers distinct from the rats’ home cages. Rats engaged in free behavior while tethered through a commutator to a Plexon data acquisition system and time-synchronized videos were recorded for each session (Plexon, Plano, TX). Noise free data from the entire recording session were analyzed using established frequency ranges from the rodent literature [delta (Δ) = 1-4 Hz, theta (θ) = 5-10 Hz, alpha (α) = 11-14 Hz, beta (β) = 15-30 Hz, low gamma (lγ) = 45-65 Hz, and high gamma (hγ) 70-90 Hz (28,29)] and standard LFP signal processing was used to characterize the power spectral densities (PSDs) within, and coherence between brain regions for each rat using custom code written for Matlab R2017b. A fourth order Chebychev type I notch filter centered at 60 Hz was applied to all of the data to account for 60 Hz line noise. The data was then down-sampled by a factor of five from 2 kHz to 400 Hz. A threshold of ± 2 mV was used to identify noise artifacts and remove data using intervals 12.5 milliseconds before and 40 seconds after the artifacts. To capture the power and coherence dynamics of the signal, we used only epochs that were at least 3 seconds long. For epochs that were longer than 3 seconds, we segmented them into 3-second sections removing the remainder to keep all of the data continuous over the same amount of time. An example trace LFP oscillation is shown in Figure 1A.

**Figure 1:**
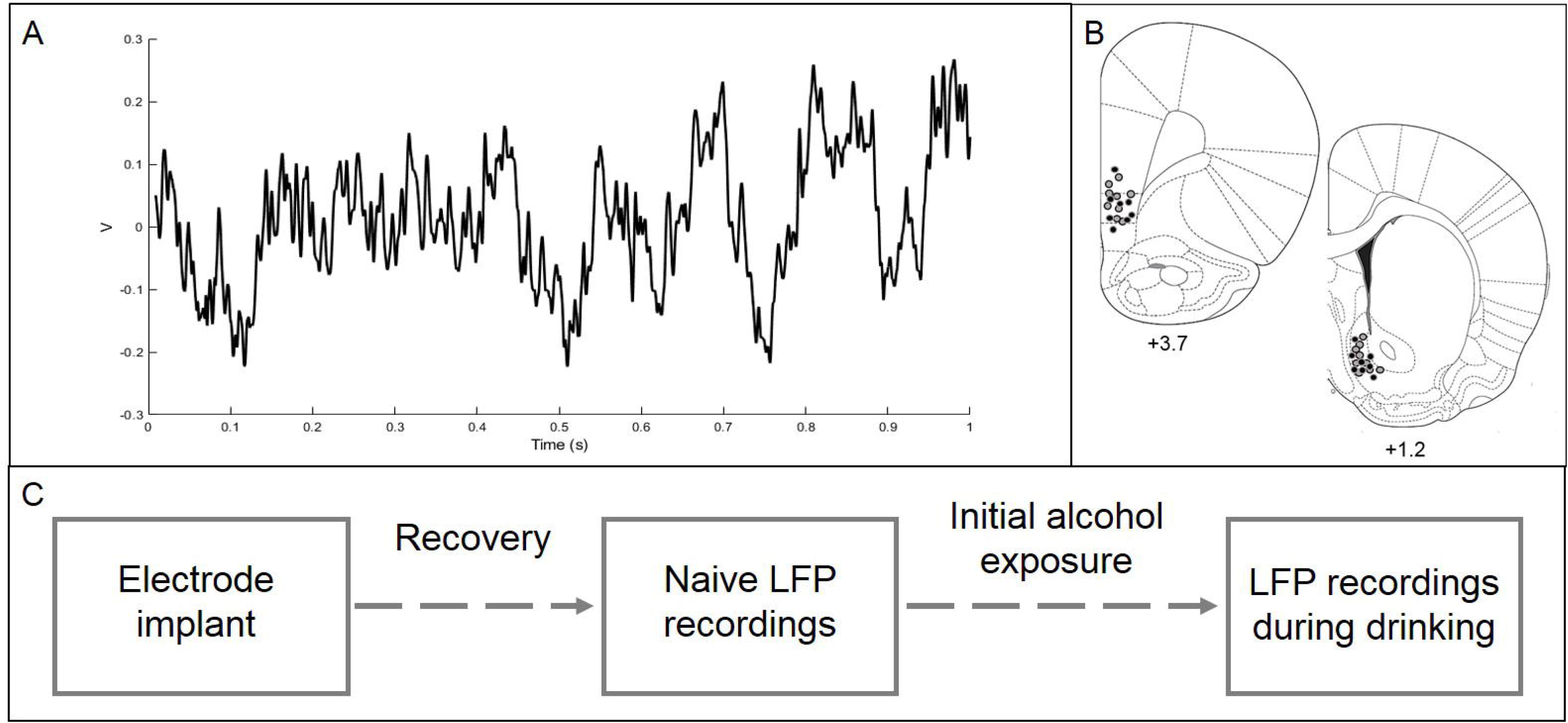
A sample trace of corticostriatal oscillations used in the prediction models (**A**). Histology figures representing electrode placements in the NAcSh and mPFC. Males are represented by black dots and females are represented by grey dots (**B**). Experimental timeline (**C**).

PSDs were computed using MATLAB’s *pwelch* function using a 1.6 second Hamming window with 50% overlap. The PSDs for each 3-second segment were then averaged together to get a single representative PSD for the 30-minute recording session. Total power (dB) was calculated for each frequency range. To account for the 60 Hz notch filter, power values of frequencies from 59 to 61 Hz were not included in the analysis. The power per frequency band was then normalized as a percent of the average total power of the signal from 1 to 90 Hz (beginning of Δ to end of hγ).

Coherence was computed using the function *mscohere* with a 1.3 second sliding Hamming window with 50% overlap. The average coherence between each pair of frequency bands from 1 to 90 Hz (excluding 59 to 61 Hz) was used to normalize the average coherence of each frequency band within that neural site pair.

### Determination of Estrous Phase

After each baseline recording session, estrous cycle was determined via vaginal lavage as described previously (13). Slides were stained using thionin and the stage of estrus was assessed using an AmScope light microscope (Irvine, CA). Proestrus was characterized as >75% of the cells in the sample being nucleated epithelial cells. Estrus was characterized as dense sheets of cornified epithelial cells, and diestrus was characterized as scattered nucleated and cornified epithelial cells, along with leukocytes (diestrus-1), or the relative lack of any cells (diestrus-2).

### Verification of Electrode Placement

At the end of the experiment, rats were euthanized using CO_2_ gas, brains were extracted and subsequently snap frozen in 2-methylbutane on dry ice. Tissue was stored at −20°C prior to being sectioned at 40 µm using a Leica CM1850 cryostat and stained with thionin. Electrode placement was verified using an AmScope light microscope (Irvine, CA). Figure 1B shows the electrode placements. Three animals’ (two males and one female) brains were not preserved properly so we were unable to verify electrode placements in those rats.

### Experimental Overview

Following one week of habituation to the animal facility, rats were implanted with bilateral recording electrodes targeting corticostriatal regions. After at least one week of recovery, baseline LFPs were recorded in two, 30-minute sessions for each male rat, and in each phase of estrous (proestrus, estrus, and diestrus) for each female rat. After baseline LFP recordings were collected, rats were allowed to drink 10% alcohol in a limited access paradigm for 9 sessions (90 minutes a day, MWF, in a neutral chamber) in order to introduce each rat to alcohol. Animal weights and the volume of alcohol consumed was measured following each session in order to calculate g/kg of alcohol consumed. Next, LFP oscillations were recorded without access to alcohol for 15-min, and then with access to alcohol for 30-min, across five distinct sessions. It is important to note that the male rats in this study were also used for a separate study investigating the impact of deep brain stimulation on alcohol drinking behaviors. See Figure 1C for an experimental timeline.

## Statistical Analysis

### Linking corticostriatal LFPs to biological sex and phase of estrous

In order to link corticostriatal activity to biological sex or phase of estrous we used an unbiased machine learning approach similar to what we have published previously (30,31). We built predictive models using corticostriatal LFPs to classify rats by biological sex and female rats by phase of estrous. Each recording session produced 60 LFP features: 24 measures of power (6 frequency bands X 4 channels) and 36 measures of coherence (6 frequency bands X 6 channel combinations). We used a penalized regression method (lasso) in order to capture potential combinations of LFP features that correlated with biological sex or phase of estrous. The Matlab package Glmnet (32) was used to implement the lasso using a 4-fold cross-validation with 100 repetitions for each of the following models: 1) male vs. female (diestrus); 2) male vs. female (estrus); 3) male vs. female (proestrus); 4) diestrus vs. estrus; 5) diestrus vs. proestrus; and 6) estrus vs. proestrus. The accuracy of the model is reported as the average cross-validated accuracy.

### Permutation testing

In order to assess the relative accuracy of the prediction models, we compared the real model performance to models built and tested on 100 different *random permutations* of the data (see Figure 2). As the outcomes of these models are binary, the random permutation models should reflect chance predictions. Thus, if the real models performed better than chance, we determined that there is some information in the circuit related to our binary outcome. However, since our sample sizes were relatively small and we used multiple recording sessions from the same animal as separate samples in the real model, we also evaluated models built on permutations of binary rat groupings (*group permutations*). This was done by keeping the LFP oscillation data together with the animal it was recorded from and shuffling the group assignment of each animal’s set of recordings. This permutation test evaluated the information contained within LFPs about all possible rat groupings. Biological sex was equally represented in each group permutation. We calculated the mean accuracy and 95% confidence intervals of cross-validated accuracy from the real, random permutation, and group permutation distributions, as well as z-scores comparing the real and random permutation distributions.

**Figure 2:**
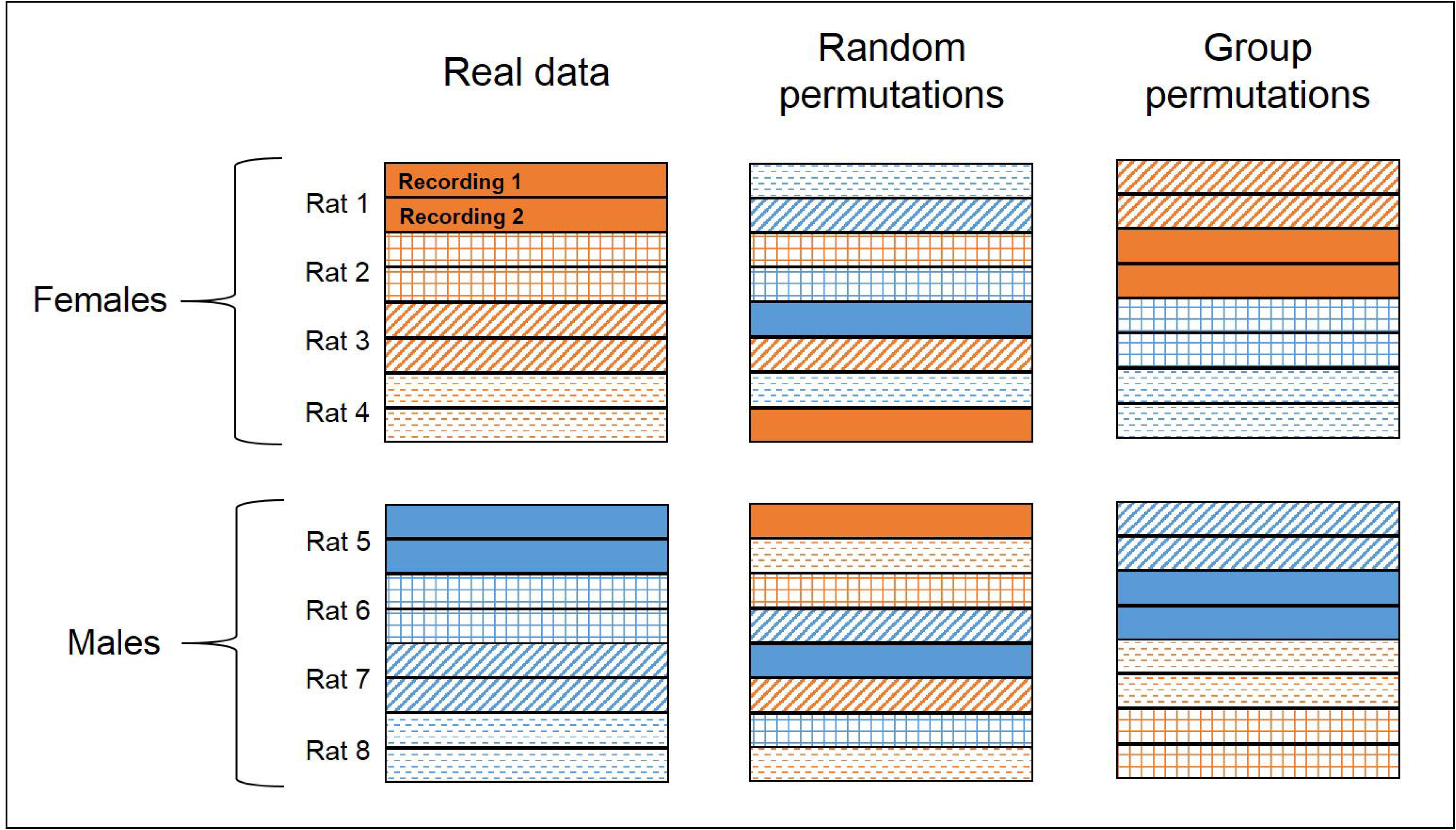
Schematic representation of the permutation testing. Each set of bars represents data from one rat (if each rat has two recordings), with males in blue and females in orange. Randomly permuted models are built on 100 iterations of shuffled data. Group permutation models are built on all possible combinations of rats assigned to each group (e.g., male or female), but the two recordings from each rat are kept together and males and females are equally represented in each permutation.

### Linking corticostriatal LFPs to alcohol intake levels

In order to analyze the impact of hormone status on alcohol intake during the recording sessions, we used a linear mixed model because two females were lacking at least one drinking day in either estrus or proestrus. Hormone status (diestrus, proestrus, estrus, or male) was used as the fixed effect, controlling for rat identification as the random effect, to predict alcohol intake during each session.

We used a similar machine learning approach (as described above) to link corticostriatal activity to alcohol intake levels, except the outcomes were continuous (g/kg of alcohol consumed by each rat across each day) rather than binary. If the lasso indicated that information existed in the LFP signal, we implemented exhaustive single feature regressions using each LFP predictor to determine the relative information content of each feature, as we have previously described in detail (31).

## Results

### The ability of corticostriatal LFPs to predict biological sex depends on female estrous phase

Models built from corticostriatal LFP features were able to outperform randomly permuted data in predicting biological sex, and the accuracy of the model performance depended on the hormone status of the females. Models predicting males vs. females in diestrus performed with the highest average accuracy; Figure 3 shows the predictive models for males vs. females in diestrus (random permutation μ = 54.96±0.6%, real μ = 71.01±1%, z = 1.71; 3A), males vs. females in proestrus (random permutation μ = 43.85±0.8%, real μ = 57.7±1.5%, z = 1.09; 3B), and males vs. females in estrus (random permutation μ = 48.15±0.6%, real μ = 56.1±1.3%, z = 0.81; 3C). It is important to note, however, that models built on group permutations of male vs. females in diestrus performed just as well as the real models (group permutation μ = 73.28±0.0002), indicating that the magnitude of sex-based differences corticostriatal circuit activity was no greater than random groupings of rats (balanced for sex) in this sample.

**Figure 3:**
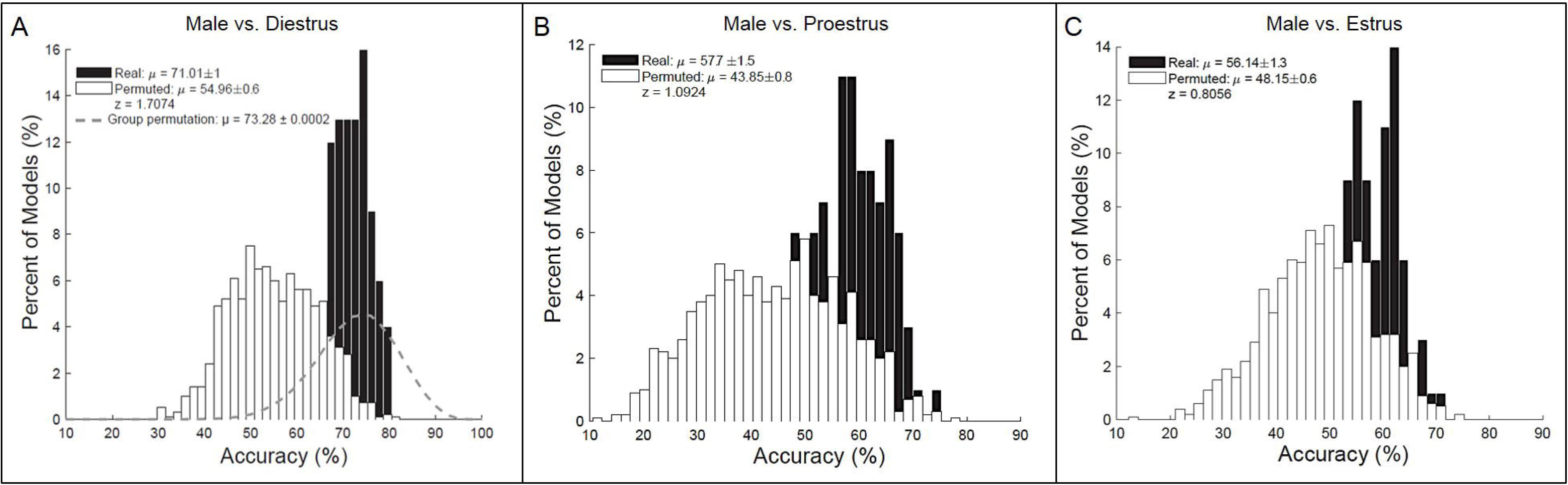
Biological sex (i.e., trait-level) prediction models (n=10/sex X 2 recordings/rat). Corticostriatal LFP oscillations predicting males vs. females in diestrus (random permutation μ = 54.96±0.6%, animal permutation μ = 73.28±0.0002%; real μ = 71.01±1%, z = 1.71; **A**), males vs. females in proestrus (random permutation μ = 43.85±0.8%, real μ = 57.7±1.5%, z = 1.09; **B**), and males vs. females in estrus (random permutation μ = 48.15±0.6%, real μ = 56.1±1.3%, z = 0.81; **C**).

For the female rats, the accuracy of models built from corticostriatal LFP features to predict phase of estrous fluctuated based on hormone status. Models predicting estrus vs. diestrus performed with the highest accuracy; Figure 4 shows the predictive models for estrus vs. diestrus (random permutation μ = 50.72±0.6%, real μ = 64.92±1.2%, z = 1.57; 4A), estrus vs. proestrus (random permutation μ = 40.97±0.6%, real μ = 53.94±1.5%, z = 1.38; 4B), and diestrus vs. proestrus (random permutation μ = 57.49±0.6%, real μ = 51.74±1.1%, z = −0.65; 4C).

**Figure 4:**
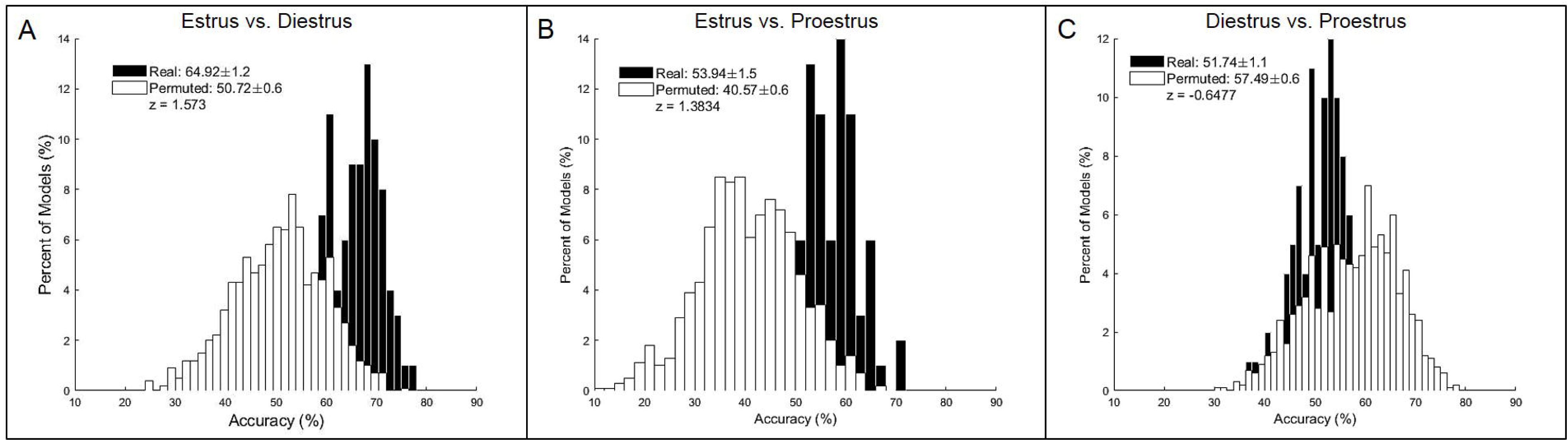
Phase of estrous prediction models (n=10 × 2 recordings/phase). Corticostriatal LFP oscillations predicting estrus vs. diestrus (random permutation μ = 50.72±0.6%, real μ = 64.92±1.2%, z = 1.57; **A**), estrus vs. proestrus (random permutation μ = 40.97±0.6%, real μ = 53.94±1.5%, z = 1.38; **B**), and diestrus vs. proestrus (random permutation μ = 57.49±0.6%, real μ = 51.74±1.1%, z = −0.65; **C**).

### Corticostriatal LFPs predict alcohol intake levels in males, but not females

Due to headcap failures, only 5 rats from each sex were able to be recorded following being trained to drink alcohol. A linear mixed effect model indicated that hormone status significantly impacted alcohol intake levels [F(3,17.32) = 4.11, p<0.05], with males drinking significantly less alcohol than females in diestrus (p = 0.052; 5A). During proestrus and estrus, female drinking amounts were not significantly different than male drinking amounts (p = 0.073 for proestrus; p = 0.702 for estrus).

We also evaluated whether we could predict biological sex in the context of alcohol drinking by using LFP oscillations collected during alcohol consumption. Figure 5B shows the predictive models for males vs. females in diestrus (random permutation μ = 44.99±0.2%, real μ = 86.81±0.01%, z = 3.76; group permutation μ = 86.55±0.0008) while alcohol was available. Again, corticostriatal oscillations do not contain more information regarding biological sex (in the context of alcohol drinking) than information about all possible groupings of rats balanced for sex.

**Figure 5:**
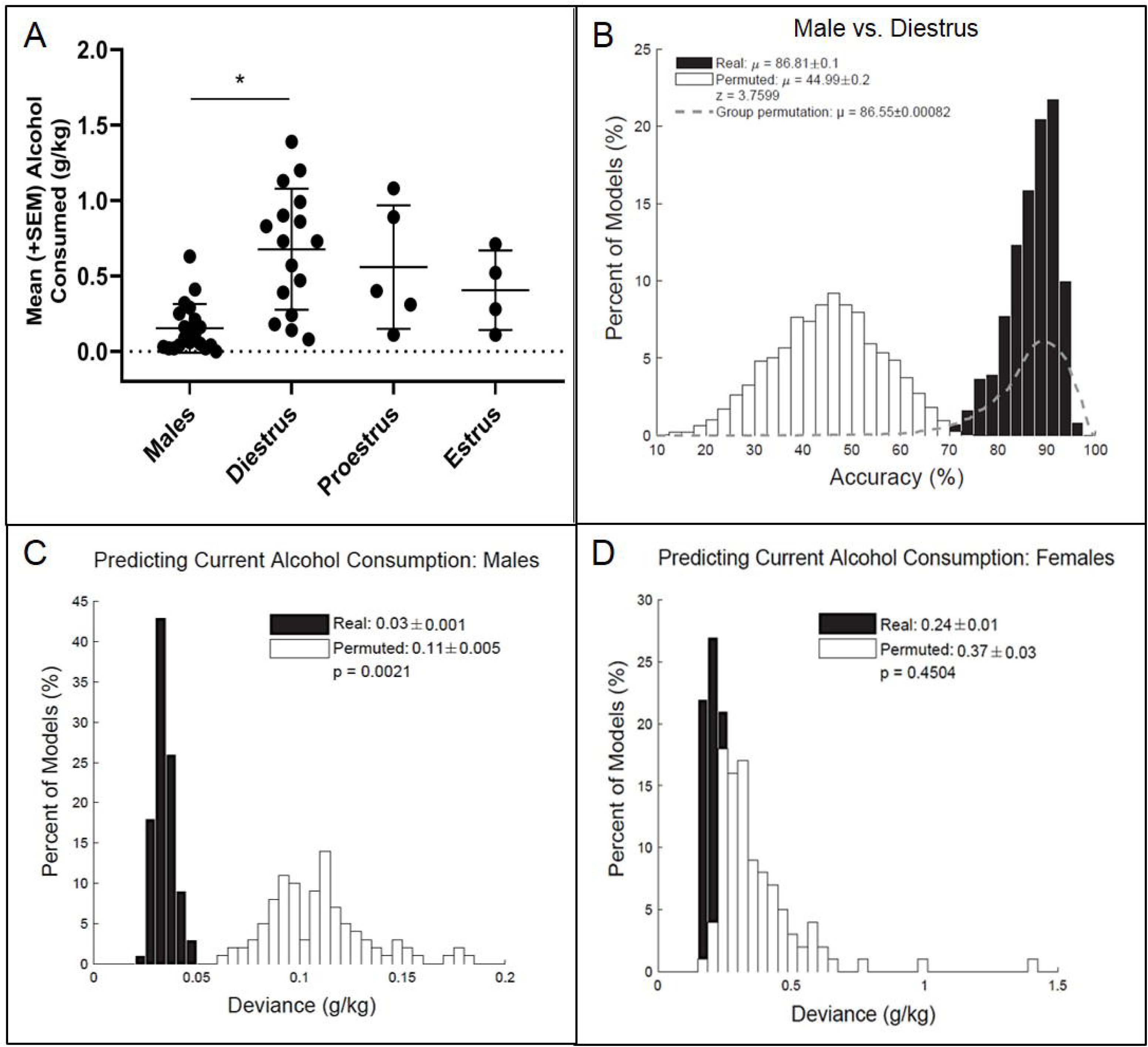
Predicting alcohol intake levels (n=5/sex X 5 recordings/rat). Female rats in diestrus drank more alcohol than male rats (p = 0.052; **A**). Corticostriatal LFP oscillations predicting males vs. females in diestrus during alcohol intake sessions (random permutation μ = 44.99±0.2%, animal permutation μ = 86.55±0.0008, real μ = 86.81±0.01%, z = 3.76; **B**). Corticostriatal LFP oscillations predict alcohol intake levels in males (random permutation error = 0.11±0.005, real error = 0.03±0.001, p < 0.01; **C**), but not in females (random permutation error = 0.37±0.03, real error = 0.24±0.01, p = 0.45; **D**).

Notably, models built from corticostriatal LFPs to predict alcohol intake levels were able to outperform randomly permuted data in males (random permutation error = 0.11±0.005, real error = 0.03±0.001, p < 0.01; 4C), but not in females (random permutation error = 0.37±0.03, real error = 0.24±0.01, p = 0.45; 4D). Table 1 lists the top five neural features important in predicting alcohol naïve males vs. females in diestrus, as well as the amount of alcohol males consumed.

**Table 1:**
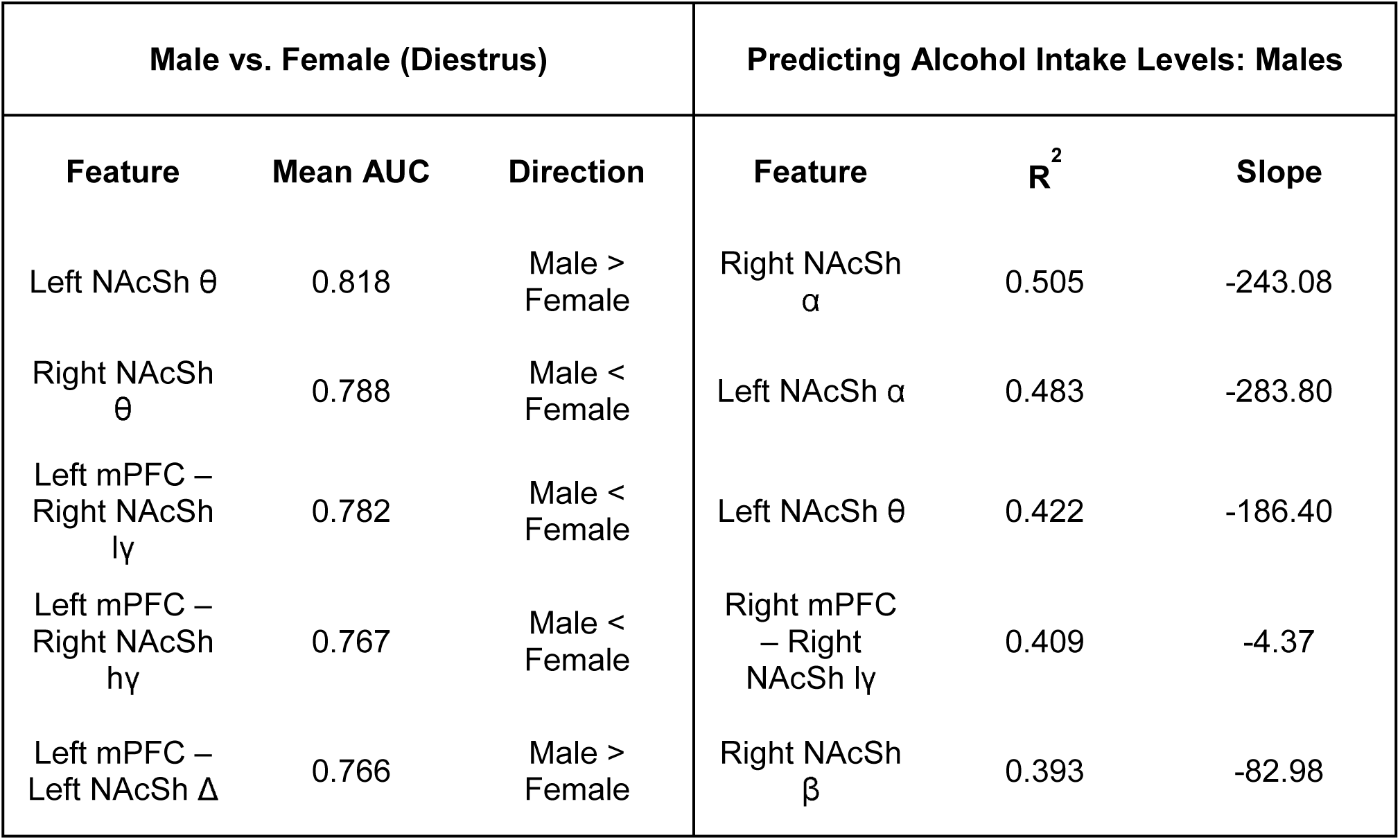
Neural features important in model prediction accuracies. The top 5 LFP features used in models predicting males vs. diestrus females and alcohol intake levels in males. Frequency bands [delta (Δ), theta (θ), alpha (α), beta (β), low gamma (lγ), and high gamma (hγ)] are described for power features within and coherence features between neural sites.

## Discussion

Here we demonstrate that LFP oscillations recorded within corticostriatal circuits contain significant information regarding alcohol intake levels in males, but not in females. We also show that while corticostriatal LFPs may contain some trait-level information (i.e., biological sex), the amount of information is similar to that observed in group permutations of animals balanced for sex. In the females, we observed small fluctuations in model accuracies as a function of ovarian hormone status, which correlated with observed differences in alcohol intake across phases of estrous and between sexes. Overall, the current experiment indicates that the neural circuits that contain information regarding alcohol consumption are sexually dimorphic.

The most compelling data from this study is that corticostriatal oscillations contained information regarding alcohol intake levels in males, but not in females. When single feature logistic regression models were applied to each neural feature, we determined that low frequency power in the NAcSh (particularly in the θ, α, and β ranges) was negatively associated with alcohol intake levels in males. Interestingly, NAcSh θ power, while negatively correlated with alcohol intake in males, also tended to be higher in males compared to females in diestrus (when males were drinking significantly lower amounts of alcohol compared to females). While these data are correlative, they do suggest that NAcSh θ power might represent a trait-level neural feature that relates to the sex differences observed in alcohol consumption. Previous studies have demonstrated that θ oscillations in the striatum, which are coherent with hippocampal rhythms, are implicated in working memory and attention tasks, and are inhibited by NAc dopamine receptor blockade (36–38). Along with the present study, these findings collectively suggest that NAc θ oscillations may be important in reward learning, and that low frequency NAcSh oscillations may perhaps serve as a potential therapeutic target in future research.

There are several potential circuits that may contain more information regarding alcohol intake levels in females. In clinical samples, women tend to use alcohol for negative reinforcement reasons, while men tend to use alcohol for positive reinforcement reasons (39). Women are also more sensitive to stress-induced relapse (5,40), and similar results have been seen in rodent models of alcohol drinking, where female rats are more sensitive to stress-induced reinstatement of alcohol seeking (41). Therefore, regions involved in emotional regulation may contain more information about female drinking behavior. One particular region of interest is the insula, which is activated by natural and drug rewards, is involved in craving, and integrates emotional stimuli contributing to mood regulation (14). Clinical studies report that reduced insular grey matter volume is correlated with increased alcohol expectancy in female problem drinkers, but not in male problem drinkers (42). Interestingly, insular activation is further enhanced by alcohol cues in alcohol-dependent women compared to non-dependent women, while men show greater alcohol cue reactivity in the striatum compared to women (43,44). In light of these previous reports, the current experiment supports the notion that different neural circuits regulate alcohol drinking behaviors in males and females.

The current findings align well with previous work describing sex differences in alcohol drinking behavior. Here we replicate findings that female rats drink more alcohol than male rats when accounting for body weight, with female alcohol intake levels fluctuating slightly across the different phases of estrous (12,13,33). Interestingly, when predicting phase of estrous in females from corticostriatal LFPs, the accuracies of the prediction models line up with differences in drinking levels across estrous phases. Specifically, the model predicting estrus from diestrus performed the best, which aligns with the phases in which female drinking behavior is most different. These data are particularly interesting considering that ovarian hormone status has been shown to influence addictive behavior in female rats and in women [though less so with alcohol and more so with other addictive substances like cocaine (5,34,35)]. Our future work will continue to investigate the role of ovarian hormones in altering substance use behaviors (and the underlying neural circuits) with the aim of developing a more comprehensive picture of the neurobiology of addiction in female rodents.

This work is further supported by previous studies using corticostriatal oscillations to characterize alcohol drinking behaviors in male rats. For instance, in male rats chronically exposed to alcohol, β power in the NAcSh is reduced during alcohol consumption periods compared to alcohol deprivation periods (21). This change in NAcSh β power coincides with an increase in NAcSh dopamine content, suggesting that changes in NAcSh β oscillations are influenced by dopamine signaling in the striatum (or vice-versa). Additionally, alcohol-preferring P rats (sex unspecified) show reduced PFC-NAc θ coherence, which is enhanced during alcohol drinking, compared to Wistar rats, suggesting that reduced connectivity in corticostriatal circuits may be related to the increased alcohol consumption in P rats (26). A significant amount of future work is required to understand the neural circuit dynamics of reward behavior across spatial resolutions (e.g., from single-cell to multi-cell to LFP recordings), but the current data supports the notion that electrical signals recorded in the NAcSh can serve as a valuable readout of substance use behaviors in male rodents.

It is important to consider one limitation to the current study. When attempting to predict males vs. females in diestrus, the real model outperformed models built on random permutations (chance), however the group permutation models had a similar accuracy to the real model. This adds a layer of complexity to the interpretation of the data, as the accuracy of the group permutations suggests that the information in the circuit regarding biological sex is no greater than the information describing natural variability in circuit activity between groups of animals (balanced for sex). There are likely many psychological domains in which corticostriatal circuit activity contains information; thus, some of the group permutations may be finding real differences between rats that are not related to biological sex. However, if biological sex was associated with substantially different corticostriatal oscillations, we would expect the real models to perform better than both the random and group permutations. It is unclear whether adding more rats to the experiment would have altered the relative accuracies of the real models and group permutations, so our future work will systematically analyze how many animals/samples are necessary to build a group permutation model with accuracies that approach chance. Nevertheless, this limitation does not reduce the importance of the present data. The models predicting alcohol intake levels in males and females were within animal, meaning the neural features identified in the continuous prediction models are directly related to the variability in alcohol intake in males. Furthermore, our future work will attempt to provide a causal link by specifically manipulating the neural features identified in the male models in the hopes of changing alcohol drinking behavior.

## Conclusions

The current dataset contributes to our long-term goal of characterizing the neural circuits that underlie alcohol drinking behavior in males and females, and our data suggest that these circuits are sexually dimorphic in nature. Moreover, the present data set reinforces the need to develop more personalized therapies for alcohol-related problems, and to help achieve this aim, current work in our laboratory aims to identify the neural circuits that underlie female alcohol drinking behavior. Additionally, we aim to characterize how circuit oscillations change across states of alcohol dependence in males and females in order to isolate (perhaps sex-specific) neural targets for reducing problematic alcohol use.

## Acknowledgments

This work was supported by funds from the Department of Psychiatry at the Geisel School of Medicine at Dartmouth (AG), the Hitchcock Foundation (AH), NIDA T32 training grant (DA037202; AH), an LRP grant from the NIH NCATS (WD), an NIAAA training grant (F31AA027441; LD), and the Dartmouth Clinical and Translational Science Institute from the NIH NCATS (KL2TR001088; WD).

